# Molecular Architecture of the Chikungunya Virus Replication Complex

**DOI:** 10.1101/2022.04.08.487651

**Authors:** Yaw Bia Tan, David Chmielewski, Michelle Cheok Yien Law, Kuo Zhang, Yu He, Muyuan Chen, Jing Jin, Dahai Luo, Wah Chiu

**Author notes:** Correspondence to: David Chmielewski, Jing Jin, Dahai Luo. These two authors contributed equally to the work (Yaw Bia Tan, David Chmielewski).

## Abstract

All positive-strand (+) RNA viruses assemble membrane-associated replication complexes (RCs) for viral RNA synthesis in virus-infected cells. However, how these multi-component RCs assemble and function in synthesizing, processing, and transporting viral RNAs to the cytosol remains poorly defined. Here, we determined both the structure of the core RNA replicase of chikungunya virus (family *Togaviridae*) at a near-atomic level and the native RC architecture in its cellular context at the subnanometer resolution, using *in vitro* reconstitution and *in situ* electron cryotomography, respectively. Within the core RNA replicase (nsP1+2+4), the viral RNA-dependent RNA polymerase nsP4, in complex with nsP2 helicase-protease, was found to co-fold with the membrane-anchored nsP1 RNA-capping dodecameric ring and is located asymmetrically within nsP1 central pore. This complex forms the minimal core RNA replicase, while the addition of a large cytoplasmic ring next to the C-terminus of nsP1 forms the holo-RNA-RC as observed at the neck of spherules formed in virus-infected cells. These results represent a major conceptual advance in elucidating the molecular mechanisms of RNA virus replication and the principles underlying the molecular architecture of RCs, likely to be shared with many pathogenic (+) RNA viruses. At last, our study will direct the needed development of antiviral therapies targeting RCs of pathogenic viruses.

**Summary:** CryoEM structure of the chikungunya virus replication complex reveals a multicomponent RNA synthesis nanomachine embedded in the plasma membrane of the host cell.

## Main Text

Chikungunya virus (CHIKV, genus *Alphavirus*, family *Togaviridae*) is a mosquito-borne pathogen that has spread globally and caused a significant health burden of acute febrile illness progressing to debilitating polyarthritis that often becomes chronic and long-lasting in millions of people. In addition, CHIKV can also induce rare but lethal encephalitis. The alphavirus (+) RNA genome is 11.8 kb in length, containing a 5’ N7-methylguanylated cap and a 3’ polyadenylated tail. It includes two coding regions: the first comprises two-thirds of the entire genome encoding for four nonstructural proteins (nsPs) in form of polyprotein precursor(s) while the second (located downstream of the subgenomic promoter) encodes for structural proteins (C-E3-E2-6K-E1) for virion assembly. Alphaviruses replicate their genomes in membrane-derived ultrastructures termed spherules that contains the negative-strand (-) RNA template likely in form of dsRNA intermediate species, and the RC assembled from nsPs (*1, 2*). The RC likely creates a favorable compartment for RNA synthesis that minimizes the host immune response to dsRNA intermediates and possesses multifunctional enzymes capable of orchestrating rapid and efficient synthesis of viral genomic and subgenomic RNAs (*3-8*). nsP1-4 are known to localize to the RCs and are essential to viral RNA replication through their own distinct enzymatic and non-enzymatic functions: nsP1 displays methyl- and guanylyltransferase activities required for viral RNA 5’ cap synthesis and plasma membrane anchoring ability; nsP2 consists of an N-terminal superfamily 1 RNA helicase and a cysteine protease at the C-terminal region responsible for polyprotein autoprocessing; nsP3 contains an ADP-ribosyl binding and hydrolase domain and a disordered region as the host factor interaction hub; lastly, nsP4 is the RNA-dependent RNA polymerase (RdRp) for viral RNA synthesis. Despite having all of the high-resolution individual structures of the nsP1-4, the overall architecture and assembly mechanisms of the alphavirus RC remains unresolved. The ultrastructure of the distantly related nodavirus Flock House virus (FHV) RC provided the first structural insight for RC assembly in the form of a dodecameric crown-shape scaffold (*7*). Similarly, multimeric ring-shaped ultrastructures were recently reported in both coronavirus RC (*8*) and alphavirus nsP1 (*9, 10*). However, these ultrastrectures have yet to explain the organization of multi-component RCs and the molecular mechanism by which RCs ultimately achieve RNA synthesis and transport to the cytosol.

Here, we applied complementary single-particle cryogenic-electron-microscopy (cryoEM) and electron-cryotomography (cryoET) methods to determine the molecular architecture of the alphavirus RC. The molecular structure of the reconstituted RC core (nsP1+2+4) fits nicely into part of the active RC density obtained from sub-tomographic averaging of the necks of the alphaviral spherules. Together, these data provide unprecedented molecular basis of the alphavirus genome replication process, likely applicable to other (+) RNA viruses such as coronaviruses and flaviviruses.

## Results

### Reconstitution of a functional alphavirus RNA replicase

To reconstitute a functional alphavirus RC, we prepared soluble recombinant nsP1, nsP2, nsP3, and nsP4 separately and ensure the enzymes are well folded and functional using previously established methods (*9, 11-13*). For this purpose, we used nsP1, nsP2, nsP3 from CHIKV, and nsP4 from o’yongnyong virus (ONNV) because we could only obtain soluble proteins of the full-length nsP4 from ONNV but not CHIKV. ONNV nsP4 (referred as nsP4 hereafter) is functionally interchangeable with the closely-related CHIKV nsP4 in the trans-replicase system (*9, 10*). Using a tandem purification strategy, the successfully reconstituted complex contains the full-length proteins of nsP1, nsP2, and nsP4 but not nsP3. Judging from the band intensity in the SDS-PAGE, nsP1 is in great excess of nsP2 and nsP4 **(Fig. 1A-B)**. To measure the biological functionality of the reconstituted complexes, we applied an RNA elongation assay to measure the polymerase activities of nsP4 alone, nsP1+4 complex, and nsP1+2+4 complex. Using a hairpin RNA template with 5’ octauridine-overhang, the recombinant alphavirus nsP4 proteins were shown to have weak *in vitro* RNA polymerase activity previously (*11*). Here, the polymerase activities of the nsP4 on the same T1 RNA template were significantly enhanced upon assembly into stable complexes of nsP1+4 and nsP1+2+4 **(Fig. 1C)**. In contrast to nsP4 alone, nsP1+4 and nsP1+2+4 catalyzed the formation of the expected full-length RNA product (RP) in 15 minutes. nsP1+2+4 complex generally outperformed nsP1+4 throughout the time course. Furthermore, the nsP1+2+4 complex was able to produce not only RP but also RP+1nt (dominant product), RP+2nts, and RP+polyA tail (faint smears above the major bands in lanes labeled nsP1+2+4) as indicated in **Fig 1C**. These results demonstrated that (1) nsP1 significantly promoted the polymerase activity of nsP4 upon forming the nsP1+4 complex; (2) nsP2 further enhanced the polymerase activity of nsP1+4 complex and boosted its terminal adenylyltransferase (TATase) activity within nsP1+2+4 complex. While nsP1+2+4 forms the core RNA replicase (referred as RC core hereafter) to catalyze the essential chemical reactions during RNA polymerization, nsP3 is likely to play additional but essential roles in viral spherule formation and virus RNA replication inside virus-infected cells.

**Figure 1.**
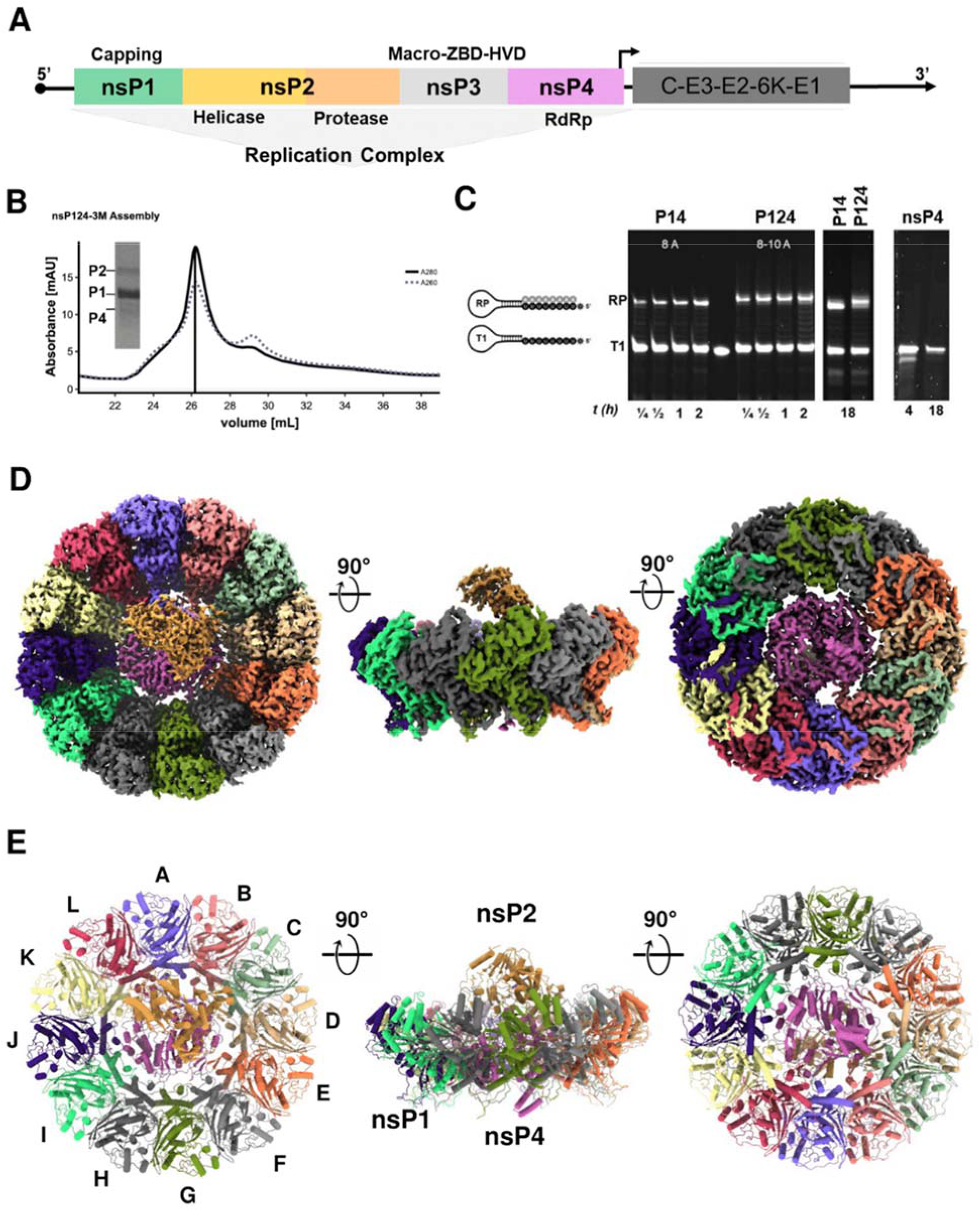
Recombinant nonstructural proteins were in vitro reconstituted via strep affinity pulldown and activated their RNA synthesis ability. **(A)** The alphavirus genome is partitioned into two open-reading frames (ORFs) where the first ORF encodes for nonstructural proteins (nsPs; nsP1+4) to assemble into a replication complex (RC) for the viral RNA replication process while the second ORF is regulated by 26S promoter (black arrow) encodes for structural proteins (SPs; C-E3-E2-6K-E1) for virion assembly. (**B)** Anion-exchange chromatography profile of RC (nsP1+2+4-3M) assembled from CHIKV nsP1 mutant (H37A; aa 1-535), CHIKV nsP2 (nsP2; aa 536-1333), ONNV nsP4 (aa 1903-2513).**(C)** The RNA polymerase activity of the RCs was observed with elongation of the T1 RNA template to its RNA products (RPs; the number of ATP incorporation labeled in white letters for each RC) at several time points (0.25-18 h) along with the nsP4 recombinant protein as a control. **(D-E)** The graphical illustrations of RC (nsP1+2+4) architecture in map representation in (D) and molecular structure in (E) at their top view, side view, and bottom view (90° rotations, from left to right). The coloring in (D-E) was assigned according to their chain numbers in the RC. (nsP1 chain coloring: violet A, salmon B, light green C, tan D, orange E, light grey F, green G, dark grey H, spring green I, indigo J, light yellow K, crimson L; nsP4 chain coloring: magenta X;nsP2 chain coloring: Peru Y; RNA of nsP2 chain coloring: green Z)

### Molecular structure of the CHIKV RC core

To gain molecular insights into the virus replication process, we determined the cryoEM structure of the nsP1+2+4 RC core at 2.8 Å resolution (**Fig. 1-2, Fig. S1-2**, and **Table S1**). The ternary complex is assembled at a stoichiometric ratio of 12:1:1, where the nsP1 dodecameric ring encloses nsP4 RNA polymerase at its central pore, and nsP2 is docked onto nsP4 of the disk-shaped nsP1+4 binary complex (**Fig. 1D-E** and **Fig. S1C**). Based on the nsP1 dodecamer structures solved recently, the crown top and the bottom skirt of nsP1 dodecameric ring face the cytoplasm and the spherule, respectively (*12, 13*). In the RC core, nsP2 extends toward the cytoplasmic side from the nsP1+4 disk (Fig. 1D-E). The total buried area between the nsP1 ring and nsP4 is 4804 Å^2^ indicating that the nsP1+4 interface is highly stable (**Fig. S3A**). Intriguingly, the nsP1 ring acts as a chaperone to co-fold with and stabilize the otherwise highly dynamic nsP4. Specifically, the nsP1 ring uses the oligomeric α-helical bundles (aa 335-364) on the inner face to hold nsP4 at the center of the dodecameric ring and recruits the flexible loops (aa 365-380, disordered in the structures of nsP1 alone and hereafter named as the hooking loop) from ten out of the twelve members of the nsP1 ring to hook up with nsP4 tightly (**Fig. 1-2** and **Fig. S2-3**). The ten unique nsP1:nsP4 interfaces are mapped to the surfaces of the N-terminal domain (NTD), Fingers, Palm and Thumb domains of nsP4, which are highly conserved across the alphaviruses (**Fig. S3D-E**). The hydrogen bonding occurs in seven of the ten nsP1:nsP4 interfaces **(Fig. 2A-J)**. We identified a putative molecular channel that is formed between nsP4 and nsP1 (Chains A and B) to connect the spherule compartment to the cytoplasm. In particular, nsP4 is oriented such that its RNA entry and exit pockets, of which are conserved among all (+) RNA virus RdRps, and its NTD are facing the spherule side of nsP1 whereas its NTP entry tunnel has full exposure to the cytoplasm (**Fig. 1-2**). In consequence, the C12 symmetry of nsP1 is disrupted upon binding to nsP4. The C terminal tail (aa 477-535) of nsP1 remains disordered, suggesting that these residues are not involved in any interaction with nsP2 or nsP4 (Fig. 2O and Fig. S4).

**Figure 2.**
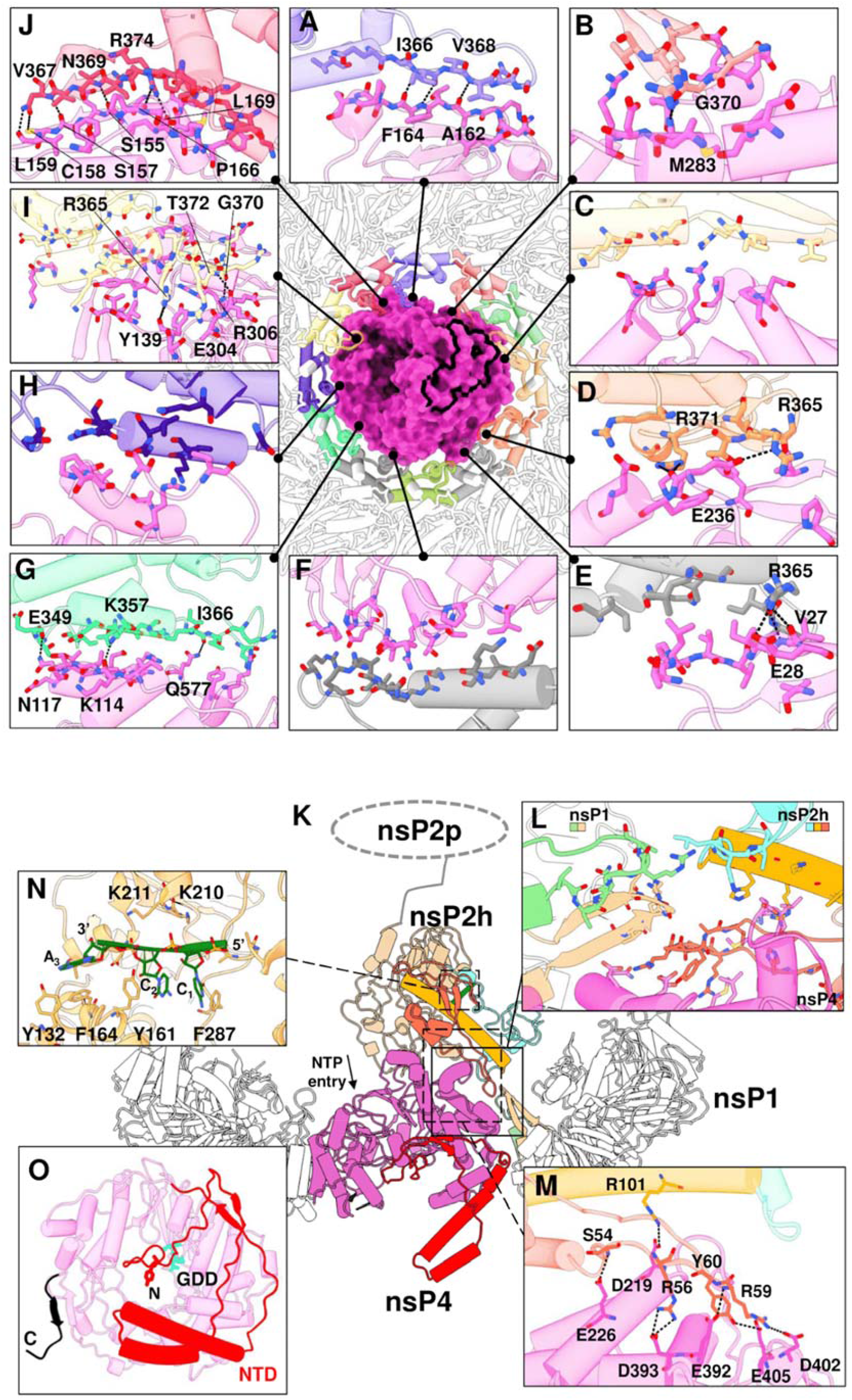
The macromolecular architecture of the RC displays a multiple interface network. **(A-J)** The interaction network of RC (made of nsP1+2+4) is presented with each colored by each subunit chain for nsP1 dodecameric ring and nsP4 (surface), based on Figure 1 chain coloring. For clarity of the interaction overview, nsP2 and its RNA ligand are hidden. Instead, the nsP2-4 interface area is outlined with a thick black line. The hydrogen bondings (black dotted lines) between the individual chain of nsP1 and nsP4 are shown here in the interface blown-up view with each residue involved named and colored according to the chain. **(K)** The overall impression of interfaces spanning across CHIKV RC (cartoon representations) where their 3-way interactions between nsP1, 2, and 4 are shown in **(L)** blown-up window (solid line and box). nsP2h (aa 1-465) region is colored according to its subdomain: NTD in orange, STALK in gold, 1B in cyan, and RecA1-A2 in peru while chain C and D of nsP1 are respectively colored in light green and tan. The unbuilt nsP2 protease (nsP2p; after aa 466) region is drawn here at the C-terminus of nsP2h for visual guidance. NTD entry site at the nsP4 motif D within the Palm (magenta region) is marked. **(M)** The hydrogen bondings (dotted lines) between nsP2-4 are listed on another blown-up window (dotted line and box). **(N)** The interacting residues from nsP2h and RNA (green) are labeled at a zoomed-in view (dotted line and box). **(O)** The bottom view of nsP4 showcases the spatial coordinates of its C-terminus (C; black; aa 600-611) and N-terminal domain (NTD; red; aa 1-105) and the active site (named GDD; cyan stick).

The interface between nsP2 and nsP1+4 subcomplex is mainly between the N-terminal NTD-Stalk region of nsP2 (helicase domain) and the Palm subdomain of nsP4 with a buried area of 906 Å^2^ (**Fig. 1-2** and **Fig. S1**,**3**). Additionally, nsP2 is close to the tips of the hooking loops of nsP1 (chains C and D), forming a three-way interaction network between nsP1, nsP2, and nsP4 (**Fig. 2L**). A flexible loop on nsP2 NTD (aa 56-65) that was disordered in the crystal structure of CHIKV nsP2 helicase (PDB 6JIM) (*14*) folds into a hairpin structure which slips into the interspace between nsP4 and the hooking loop of nsP1 (Chain C and D) (**Fig. 2L** and **Fig. S3**). Conserved residues R56, R59, Y60, and R101 from this hairpin structure established several H-bonds with D219, E392, D393, D402, and E405 from nsP4 with proximal contacts to V368, N369, and G370 from nsP1 (Chain D) (**Fig. 2L)**. While the interface between nsP2 and nsP1+4 is well resolved in the cryoEM map, the rest of the nsP2, including RecA-like domains 1-2 of the helicase core (nsP2h) and the C terminal protease region, becomes more flexible and gradually invisible (**Fig. 2K** and **Fig. S1**). Nonetheless, we identified and built a short single-stranded RNA (ssRNA) of three nucleotides within the nsP2h RNA binding groove, in a similar manner as its reported crystal structure (**Fig. 2N** and **Fig. S2**) (*14*). The bound RNA may represent the product RNA being exported from the viral spherule, before being transported to the ribosome for translation or to the capsid proteins for virion assembly.

nsP4 RdRp resides at the center of the RC core, well stabilized by the surrounding nsP1 and nsP2. Indeed, almost all 611 residues of the nsP4 are well resolved in the cryoEM density map, which includes the intrinsically disordered NTD (aa 1-105) and those disordered regions from the Fingers domain (subdivided into the index, middle, ring, and pinky fingers) not captured in the crystal structures of the alphavirus nsP4 RdRps from Ross River virus (RRV; PDB 7F0S) and Sindbis virus (SINV; PDB 7VB5) **(Fig. 1D-E, 2O and Fig. S2, S4)** (*11*). The NTD of nsP4 is composed of several characteristic structural elements (**Fig. 2O** and **S4**): (1) The N terminal tip (aa 1-25) starts from the alphavirus-conserved Y1 residue and is inserted into the central RNA binding groove; (2) One helix-turn-helix motif (aa 42-84) interacts with the pinky fingers motif and extending towards the spherule side; (3) Anti-parallel β-strands (aa 26-30 and 98-102) along with the surrounding unstructured loops wrap around the RdRp core domain. The C terminal tail of nsP4 is also well-folded and points to the spherule side of the RC core. Overall, the well-folded nsP4 structure within the RC core explains the enhanced polymerase activities (**Fig. 1C**) and reveals both unique and conserved molecular features when compared to the structurally-related RdRps from other (+) RNA viruses (**Fig. S4**).

### Molecular organization of CHIKV RC *in situ*

To validate the biological relevance of the *in vitro* reconstituted alphavirus RC core and understand its functions in the native RC in the virus-infected cells, we performed cryoET of CHIKV-infected cells at 8 hours post-infection, a time point when all RCs should be mature with fully-processed nsPs and dsRNA replication intermediate (*15, 16*). Unlike spherules of other alphaviruses that form at the plasma membrane (PM) and are subsequently internalized to cytopathic vacuoles via PI3K-Akt activation, CHIKV spherules stay at the PM without internalization (*17*). Taking advantage of the predominant location of CHIKV spherules at the PM, we collected tomograms of the periphery of CHIKV-infected cells decorated with many spherules filled with ball-of-yarn-like densities, likely corresponding to dsRNA intermediates. CHIKV spherules bulge outward from the PM with membrane-associated macromolecular complexes, assumed to be active RCs, gating at the spherule neck (**Fig. 3A-B**). Spherules were generally balloon-shaped with an average short and long axis of 63.1 ± 4.7nm and 79.0 ± 6.5nm (N=101 spherules), respectively. Below the spherules, dense condensates in the cytosol were often observed proximal to the neck (**Fig. 3C**). Such density can be attributed to RNA, RNA/Capsid mixture, and/or cell host factors recruited to active replication sites (*3, 18, 19*).

**Figure 3.**
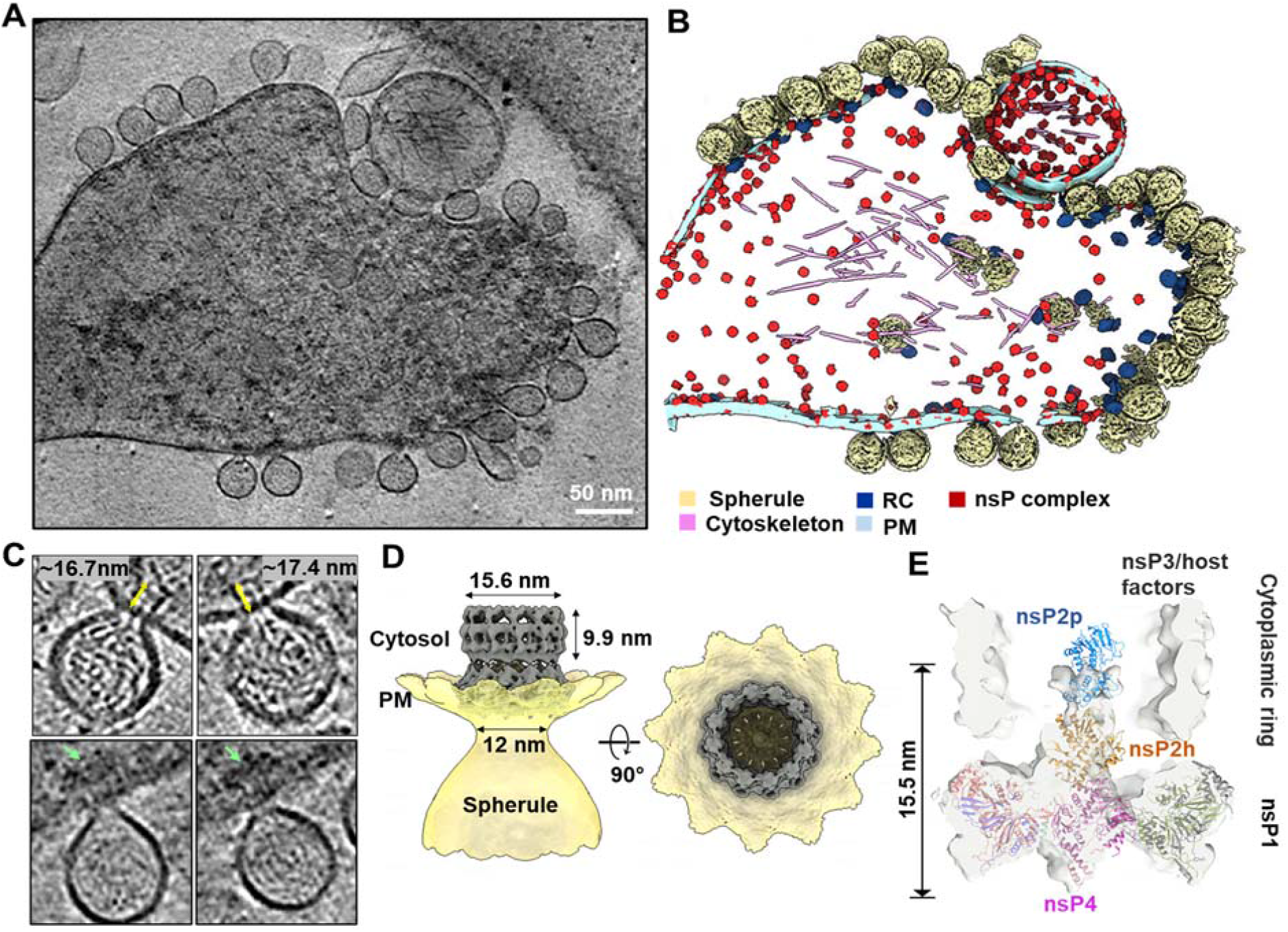
CHIKV RNA replication spherule structures revealed by cryoET. **(A)** Tomographic slice of cell periphery depicting CHIKV RNA replication spherules at the plasma membrane. **(B)**Corresponding 3D segmentation of cellular features. See also movie S1. Scale bar 50 nm. **(C)** Snapshot of the individual spherules. Yellow arrows measure the ordered density within the center of the cytoplasmic ring. Green arrows mark the additional associated density below the RC cytoplasmic ring. **(D)** CHIKV spherule 3D volume is determined by subtomogram averaging with imposed C12 symmetry. **(E)** The replicase core complex is fitted into the C1 subtomogram average map of the RC. A cytoplasmic ring as observed in **(C)**, likely made of nsP3, RNA, and host factors remain loosely connected to the nsP1 ring which is bound at the neck of the spherule. The extra density above the nsP2h region is likely to be the C terminal protease of nsP2, stabilized or restrained by the cytoplasmic ring.

Subtomogram averaging of the spherule neck regions revealed a 12-fold symmetric complex at 7.3 Å resolution, consisting of a membrane-associated crown ring connected to a larger, smooth ring on the cytoplasmic side **(Fig. 3D, Fig. S5)**. The membrane-associated crown ring closely matches the reported cryoEM structures of the recombinant nsP1 dodecamer (*9, 10*). While the crown ring contains a plug-like density in the 12-fold symmetric average, the cytoplasmic ring possesses a notably wide and mostly empty cavity (15.6 nm diameter, ∼10 nm long) (**Fig. 3D**). Together, the RC extends approximately 17 nm from the bottom of the crown ring to the top of the cytoplasmic ring (**Fig. 3C**). In tomogram slices, contiguous linear density was observed, which we attributed to RNA transported from the spherule through the crown and the center of the cytoplasmic ring. Such density was blurred out in the average by the imposed 12-fold symmetry. Interestingly, the two rings in the subtomogram average are linked with a fixed orientation by thin density, potentially nsP1 C-terminal helices that are unresolved in the nsP1 dodecamer cryoEM structures (**Fig. S5**) (*12, 13*). Future work is warranted to determine how the two rings assemble into the active RC and the other cofactors that potentially guide their assembly.

We then determined an asymmetric reconstruction of the RC, revealing that the plug density in the center of the crown is positioned asymmetrically with a small gap on one side (**Fig. 3E, Fig. S5)**. This is consistent with the structure of the active nsP4 RdRp in the RC core complex that contacts 10 of 12 nsP1 hooking loops (**Fig. 1-2**). Above the crown, significant density was resolved in the center of the cytoplasmic ring. The abovementioned RC core complex (**Fig. 1D-E**) docked nicely into this cryoET reconstruction, while additional density above the nsP2h domain is likely the protease domain (nsP2p) that is unresolved in the reconstituted RC core complex (**Fig. 3E**). It is possible that the cytoplasmic ring chamber confines this highly flexible region and/or nsP2p forms loose interactions with RNA and/or the interior of the ring that stabilizes its conformation in the mature RC. Since nsP3 is missing from the reconstituted RC core complex and known to localize to the active RC, we postulate the cytoplasmic ring in our reconstruction comprises nsP3, newly synthesized viral RNA, and possible host factors. In total, these reconstructions reveal both the composition of the viral RC core in the native cellular context and the overall molecular architecture of this elegant nanomachine for viral RNA production.

### Excess nsPs form membrane-associated nonreplicative complexes

In addition to the RCs positioned at the neck of spherules, we observed a large number of ring-like complexes smaller than RCs docked to the inner leaflet of the PM in the absence of RNA or membrane invaginations. Interestingly, these macromolecular complexes were especially numerous on thin cell extensions with underlying bundled cytoskeleton filaments (**Fig. 4A-B**, particles in red named as nsP complex), reminiscent of the reported distribution of nsP1 in filopodia (*20, 21*). Occasionally, we also observed extracellular membrane vesicles of various sizes that were decorated with such complexes on the inner leaflet (**Fig. 3A-B**). To reveal the molecular identity of the complexes, we determined a subtomogram average of over 5,000 extracted particles to ∼10 Å resolution **(Fig. S5)**. The reconstruction revealed a 12-fold crown ring ∼20 nm in diameter with a 7 nm central pore that closely matches the nsP1 dodecamer reported (**Fig. 4C**) (*12, 13*). We then considered whether these rings were nsP1 alone or comprised multiple assembly states of nsPs. To answer this, we determined an asymmetric reconstruction of the rings, revealing asymmetric density plugging the center of the crown and some density again above it on the cytoplasmic side. This density corresponded nicely to the RC core complex described above (**Fig. 4D, Fig. 1C-D**). Noticeably, the nsP2 attributed density only included the helicase domain, while the protease region and the large cytoplasmic ring of the RC were not detected. To determine whether there were any nsP1 rings alone, we performed 3D classification on this asymmetric complex, which revealed all the rings were composed of RC core members: nsP1+2+4 (**Fig. S6C**). Therefore, while expression of nsP1 alone is apparently responsible for the induction of thin extensions (*20, 21*), it is likely achieved via the RC core complex formations in the virus-infected cells although we cannot exclude that nsP1 functions in a form not detected in our study. Since our time point of imaging (8 h.p.i) is considered late in the CHIKV replication cycle, the production of large amounts of cleaved nsPs likely results in extreme molar excess of the nsP1+2+4 complex relative to template strand RNAs. Therefore, we consider such complexes nonreplicative due to the absence of the cytoplasmic ring and the spherule (and template RNA).

**Figure 4.**
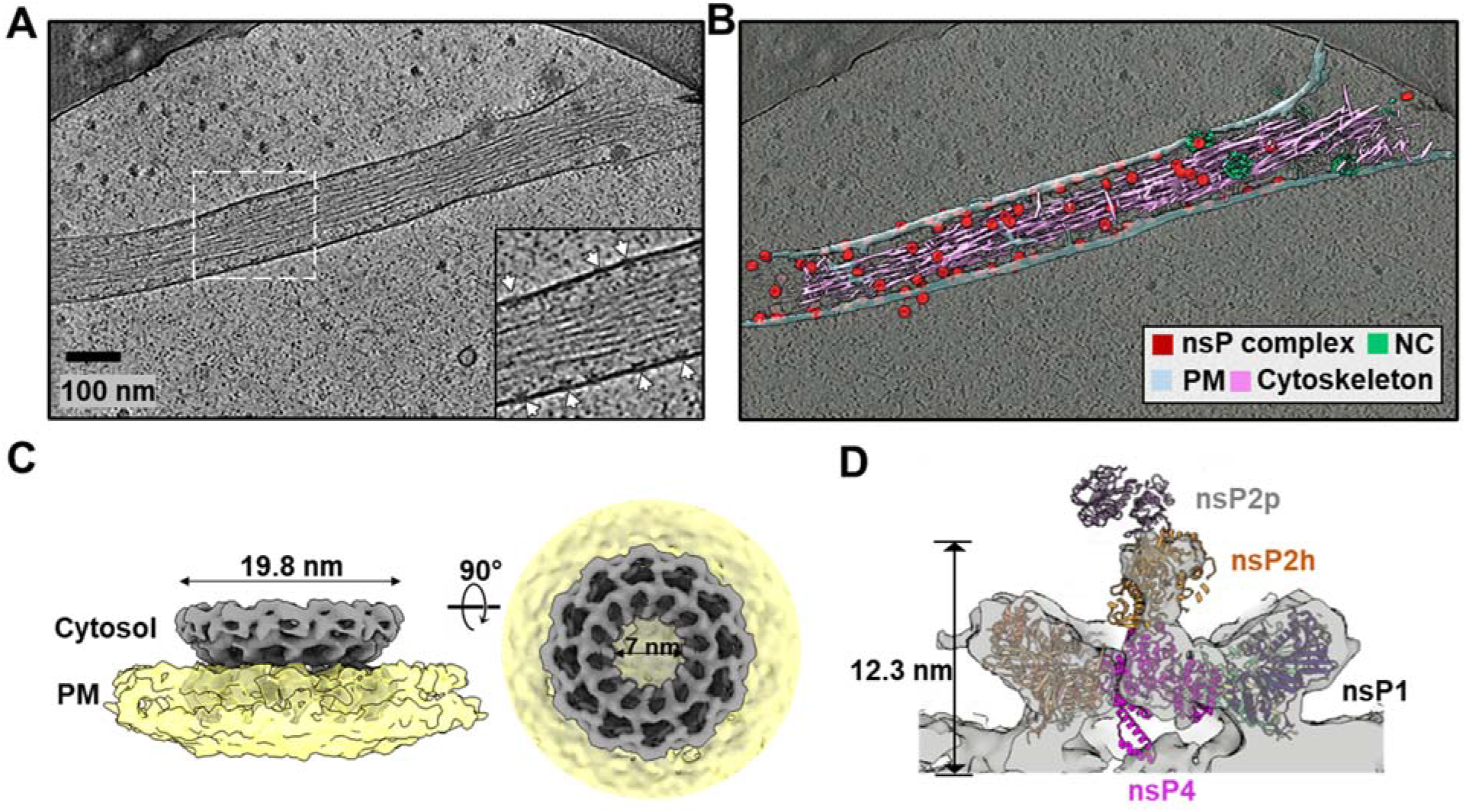
Inactive nsP complex induced membrane protrusion revealed by cryoET. **(A)** Tomographic slice of cell periphery depicting a filopodia-like membrane protrusion structure extended from the plasma membrane. White arrows point to the membrane-associated nsP complexes. **(B)** Corresponding 3D segmentation of cellular features. See also movie S2. Scale bar 100 nm. **(C)** CHIKV nsP complex 3D volume determined by subtomogram averaging with imposed C12 symmetry. **(D)** The replicase core complex is fitted into the C1 subtomogram average map of the nsP complex. Note that the cytoplasmic ring as observed in Figure 3. (**C)** is absent in this inactive nsP complex. Consequently, no map coverage of the C terminal protease of nsP2.

## Discussion

In this study, we performed a multiscale study for the structural and functional characterization of the CHIKV replication machinery *in vitro* and in the infected cell using combinatory cryoEM methods. Our work provides a comprehensive view of how the alphavirus nsPs assemble into a RCs, that recruit viral RNA and additional viral and host cofactors, and construct the mature viral RNA production factory – spherule on the PM.

Viral RdRps play a central role in the virus replication process and serve as a top target for antiviral development (*22, 23*). Unlike other viral RdRps from the *Piconaviridae (24)* and *Flaviviridae (25)* that possess relatively good *in vitro* RNA polymerase activity, we showed that alphavirus nsP4 gets activated upon proper folding within the central pore of nsP1 dodecameric ring. nsP2 binding further unlocked the TATase activity which is essential for the 3’-end polyadenylation of the genomic and subgenomic RNAs (**Fig. 1**). This is functionally similar to the coronavirus nsp12 RdRp which requires cofactors nsp7 and 8 for proper binding to the RNA substrate to gain decent polymerase activity (*26-28*). Based on this nsp4 localization, it is intriguing to speculate that density in the center of the coronavirus nsp3 hexamer, unresolved in the subtomogram averages with applied rotational symmetry, represents the coronavirus RdRp (Fig. S8) (*29*). This conserved RC architecture would provide an elegant solution to the mechanisms of RNA synthesis, immediate capping of nascent RNAs, and sealing the membrane microcompartment while exporting RNA to the cytosol for translation and packaging into virions.

The NTD subdomain of nsP4 is unique with its N-terminus inserted into the RNA binding groove, which is equivalent to the priming loop of dengue NS5 RdRp (*25, 30, 31*) (**Fig. 2 and Fig. S4**). As a result, the N-terminus of nsP4 NTD needs to move away to allow the binding of the dsRNA elongation substrate (*32*). The helix-turn-helix motif in nsP4 NTD that protruded into the spherule lumen may also provide an additional binding footprint for supporting dsRNA. The putative molecular channel between nsP4 and nsP1 (Chains A and B) may serve as a bidirectional ssRNA translocation channel to export RNA products to the cytosol as well as to import the parental viral (+) RNA for (-) RNA synthesis **(Fig. 5)**. The alphavirus-unique nsP4 NTD could also evolve adaptively to organize the flexible loops of NTD and Fingers subdomains into an extensive surface for enabling nsP1:nsP4 interactions. Only one copy of nsP2 is found to bind to nsP4 through its NTD region at the cytoplasmic side. This molecule of nsP2 improves the polymerase activity on the short RNA substrate, likely via binding and stabilizing the active conformation of nsP4 (**Fig. 1-2**). As a Superfamily 1 RNA helicase, nsP2h translocates along ssRNA in a 5’>3’ direction (*33*). Given its relative location and orientation to the nsP4 polymerase, nsP2h may function by pulling out the product RNA from the spherule analogous to a pulley system, as a hypothesis that requires further study.

**Figure 5.**
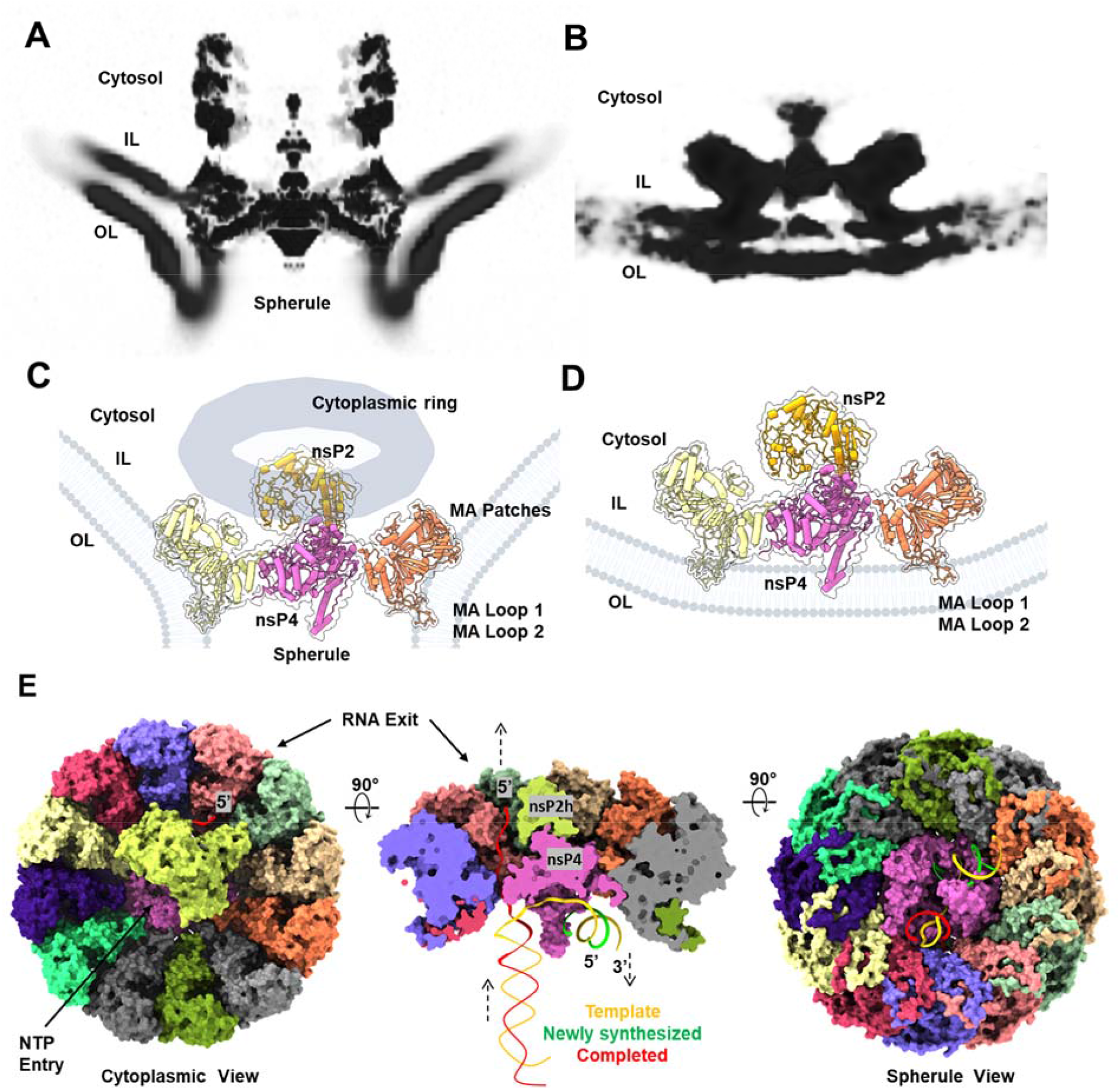
Remodeling of the host membrane is coupled to assembly of active viral RC. **(A)** A central slice view of the tomography volume at the active viral RC spherule highlights the curved plasma membrane at the nsP1 contact sites (membrane association loops 1 and 2 (MA Loops 1 and 2) from the bottom of the nsP1 ring and newly identified MA patches from the nsP1 upper ring) as shown in **(C)**. In contrast, the **(B)** panel shows the central slice view of the tomography volume at the inactive viral nsP complex which is simply docked to the inner leaflet of the plasma membrane, using the ML Loops 1 and 2 on nsP1 ring **(D). (E)** Molecular model of how the viral RC core complex facilitates viral RNA replication. the dsRNA replication fork is modeled and colored. Template RNA, ie. Antigenomic (-) RNA is in yellow, newly synthesized progeny RNA (or sgRNA transcript) is in green, and the RNA in red is the completed product RNA to be exported to the cytoplasm for translation and virion assembly. NTP entry tunnel on nsP4 polymerase and the putative RNA exit tunnel are labeled.

While nsP1 expression alone was reported to drive cell morphological change and induce the formation of thin extensions, the molecular organization of nsP1 and its mechanism of action in virus-infected cells is unclear (*34-36*). Here, we demonstrated that at the molecular level, nsP1 rings mainly in the nonreplicative form of nsP1+2+4 RC core complex are present in extremely high concentrations in thin extensions, with each inducing a small and local positive curvature on the membrane (**Fig. 4A-C** and **Fig. 5B,D**). It is conceivable that these aggregates of nsP1-induced membrane deformations stabilize high membrane curvature to drive formations of thin extensions (*21, 35*). How nsP1 induces actin remodeling remains to be determined. One interesting observation in this study is that these nsP1+2+4 RC core complexes seal the active RNA replication spherules only when they assemble with cytoplasmic rings into active holo-RCs. Unlike RC core alone docking at the PM, the RC core in the holo-RC supports a high negative curvature at the spherule neck via extensive binding of nsP1 to the membrane (**Fig. 5A-B**). We identified additional membrane association (MA) patches on the pheripheral side of the nsP1 ring (**Fig. 3E, Fig. 5A,C**, and **Fig S8**). Together, these MA loops and the MA patches around the nsP1 ring maintain a tight engagement with the negatively bent PM (**Fig. 5A,C** and **Fig. S7**), which seem to be functionally equivalent to the transmembrane domains of the nsp3 from coronavirus and the N terminal mitochondrial membrane localization sequence of FHV protein A (**Fig. S8**) (*29, 37*). It is not known if the stable yet inactive nsP1+2+4 RC core complexes may be reactivated by recruiting the nsP3-cytoplasmic ring with viral RNA and potentially replace those functional RC prone to degradation by host response. Future work is warranted to address these questions.

Based on the current knowledge, we generated a model for CHIKV RNA replication (**Fig. 5E**). The spherules are thought to contain viral dsRNA intermediates, comprising the (-) RNA template and (+) nascent RNA, and export of replicated/transcripted ssRNA species to the cytosol for translation or virion packaging. During viral RNA replication, the synthesis of a nascent RNA product and the export of the completed product RNA from the last replication are coupled and thus reaching a equilibrium state with consistent amount of RNA maintaining the 3D volume of a mature spherule. Capping the 5’-end of the product RNAs can be conveniently catalyzed by nsP2 and nsP1 right after they are exported to cytosolic surface of RC. The cytoplasmic ring of the RC that is attached above its RC core complex seems to be essential in maintaining the continuous material exchange between the spherule and the host cell cytoplasm (**Fig. 3 C-E** and **Fig. 5A,C**). Overall, the CHIKV RC structure provides the molecular basis of viral RNA replication and serves as a useful tool for antiviral development against alphaviruses and other (+) RNA viruses.

## Acknowledgments

This research is supported by the Singapore Ministry of Education under its Singapore Ministry of Education Academic Research Fund Tier 2 (T2EP30220-0020) and Education Academic Research Fund Tier 1 (2021-T1-002-021) to D.L. This research was supported by the NIH grants R01AI148382, P01AI120943 and S10OD021600 (to W.C.). We thank Dr. Andres Merits for providing the CHIKV and ONNV cDNA and insightful discussions. We thank the scientific facility support from NTU Institute of Structural Biology and Protein Product Platform. We thank SLAC National Accelerator Laboratory for access and support of these studies, and all SLAC cryoEM staff for technical support and assistance. We thank members of the DL and WC lab for their support.

## Author Contributions

YBT, DC, JJ, WC, and DL designed the study; all authors performed the experiments and analyzed the data; YBT, DC, JJ, WC, and DL wrote the manuscript with inputs from all authors.

## Data And Code Availability

The SPA-cryoEM density map of the CHIKV RC core complex nsP1+2+4 has been deposited in EM Database under the accession code EMD-XXXXX. The corresponding atomic coordinates have been deposited in the Protein Data Bank under accession code XXXX. Tomography data EMDB-XXXX.

## Supplementary Materials

Materials and Methods

Figs. S1-S?8

Table S1

References

