## Supplementary File for "Molecular Architecture of the Chikungunya Virus Replication Complex"

**This PDF file includes:**

Materials and Methods
Figs. S1 to S9
Tables S1

### Materials and Methods

#### *Recombinant protein productions of alphavirus nsPs*

The preparation of nsP1 was adapted from (*1*). The gene cassette of CHIKV nsP1 (aa 1-535) comprises affinity fusion tags of Strep-3XFLAG after A516 residue and mutation of H37A, created using In-fusion Snap Assembly kit (Takara Bio) and subcloned into pEGFP-C1 (Clontech) with eGFP eliminated. This nsP1 plasmid was expressed transiently in Expi293 expression system (ThermoFisher Scientific) by PEI MAX® (Polysciences) transfection and harvested at 4000 x g after five days. The protein expression was boosted by 10 mM sodium butyrate. The cell pellet was subsequently lysed in lysis buffer WI (50mM HEPES pH 8.0, 150mM NaCl, and 0.5mM TCEP) with 1mM PMSF protease inhibitor (Sigma Aldrich), 1% n-dodecyl-β-D-maltoside (DDM; Exceedbio), and 70 mU/ml of BioLock solution (IBA Lifesciences) on rotator at 4°C for 1.5 hours (h) and pulse-sonicated for 5 minutes. The lysate was clarified with 100,000 x g centrifugation at 4°C in XPN-100 ultracentrifuge (Beckman Coulter). The Strep-Tactin® Sepharose® slurry beads (IBA Lifesciences; named as strep-bead hereafter) were incubated in Econo-Pac® column (Bio-Rad) with the clarified supernatant. The nsP1-bound strep-bead was equilibrated to assembly buffer WII (50 mM HEPES, 150 mM NaCl, 2.5 mM $MgCl_2$, 0.5 mM TCEP, 0.008% GDN, 5% glycerol, 2.5% sucrose, ~pH 8) in 5 column volume (CV) wash for downstream RC assemblies and RNA polymerase assay. The recombinant proteins of full-length nsP2 (aa 536-1333) and its helicase domain (nsP2h; aa 536-1000) were prepared as described in (*2*). nsP3M gene (aa 1334-1659) was cloned from the same CHIKV cDNA described in (*1*) into pSUMO-LIC expression vector with N-$His_6$-SUMO tag. The expression for nsP3M was done in Rosetta II T1R *Escherichia* coli (*E.* coli) strain and cultured Luria-broth (LB) Miller media supplemented with 2.5% glycerol and 0.5 mM IPTG after induction at measured optical density (OD600) of ~0.8 for 18 h at 18˚C. The purification process for nsP3M was maintained throughout at 4˚C. The purification started with harvesting the bacterial pellet at 4,000 RPM followed by lysis with buffer A (2X PBS adjusted to 500 mM NaCl, 10% glycerol, 5 mM β-mercaptoethanol) supplemented with 1 mM PMSF in PANDAPLUS2000 (GEA) homogenizer. The supernatant obtained from 100,000 x g ultracentrifugation was incubated with Ni-NTA slurry beads (Bio Basic) for 1 h. The nsP3M-bound Ni-NTA was washed 20 CV and eluted in buffer B (buffer A plus 300 mM Imidazole). The eluant was purified subsequently in HiLoad® 16/600 Superdex 200 pg column (Cytiva) and concentrated to desired concentration to flash freeze for long-term storage at -80C till utilizations. The preparation of full-length nsP4 was adapted from (*3*). The nsP4 gene was sourced from o'nyong'yong virus (ONNV; AF079456.1; a gift from Andres Merits) and cloned into pSUMO-LIC with N-terminal hexa-histidine tag fused with cleavable SUMO tag (N-$His_6$-SUMO) for bacterial expression and purification as described in (*3*).

#### *RNA polymerase assay*

The RNA polymerase assay is optimized and uses the same T1 RNA template from (*3*). All assay reactions were performed with 1 µM protein (nsP1-4, nsP1-2-4 and nsP4) and 50 nM T1 RNA template in 0.5× assay buffer (1× buffer: 25 mM HEPES, 75 mM NaCl, 20 mM KCl, 5 mM MgAcetate, 2.5 mM MnCl2, 5 mM TCEP, 0.01% GDN, 0.5 U/µL NEB Murine RNase Inhibitor, ~pH8) prepared in nuclease-free water (HyClone). The reactions were initiated with 2 mM ATP and quenched at discontinuous time point manners for 0.25-18 h after incubating at ~25˚C in dark. The reactions were quenched with equal volume of denaturing RNA loading dye (95% formamide, 0.02% SDS, 0.01% xylene cyanol, 0.02% bromophenol blue, and 1 mM EDTA) to run on 17.5% denaturing urea-PAGE at 220 V for 1.5-2 h. The urea-PAGE was imaged via Bio-Rad Gel-Doc XR+ Imager using Fluorescence Blot ALEX488 for analysis.

***In-vitro transcription RNA synthesis***

T2 RNA was designed to have a central double-stranded region of 50 base-pairing and flanking 5’ overhangs. To synthesize T2 RNA with *in vitro* transcription (IVT), a DNA template with T7 promoter (5’ TAATACGACTCACTATA 3’) was ordered from Integrated DNA Technologies to conduct IVT with manufacturer recommendation from Hiscribe™ T7 High Yield Synthesis Kit (NEB). The IVT product was purified via phenol-chloroform extraction method and Monarch RNA clean-up Kits (NEB).

*T2 RNA sequences:*

*Sense strand:*

*5’GGGAUAAUGCAAGCAUACCGAUCUUCCAACGUUUCUCCGAACCCACAGGGACGUAGGAGAUGUUAUUUUGUUUUUAAUAUUUC3’*

*Antisense strand:*

*5’GGGAUAAUGCAAGCAUACCGAUCUUCCAACGUUGAAAUAUUAAAAACAAAAUAACAUCUCCUACGUCCCUGUGGGUUCGGAGA3’*

***In vitro reconstitution for RC assemblies***

All *in vitro* reconstitutions and sample preparations were done at 4˚C unless otherwise mentioned. The nsP1-immobilised strep-bead was incubated with the desired combinations of nsPs (nsP2, nsP3, and nsP4) to produce nsP1-2-4. The strep-beads consisting of RCs (nsP1-2-4) were eluted in the buffer WII supplemented with 10mM desthiobiotin at ~pH 8 and loaded directly to Hitrap Q-FF column (Cytiva) for linear elution up to 1 M NaCl. The eluants were concentrated at 3000-7000 x g in Amicon Ultra (Merck) and/or Pierce™ (ThermoFisher Scientific) protein concentrators of 100 kDa MWCO for RNA polymerase assay (refer below). The RCs assembled at affinity pulldown and anion-exchange chromatography were fractionated and sampled for negative staining on carbon-coated copper 300 mesh grids (EMS) for screening and optimization of RCs.

***Single-particle Cryo-EM sample preparation***

The nsP1-2-4 RC was mixed with T2 RNA and NTP at 1:10:100 molar ratio for 1 hour at ~25˚C before concentrating down to ~30 µL. The cryo-grid preparation (including graphene grid fabrication) was adapted from (*1, 4*). About 5-6 µL of the sample (estimated at ~0.2-0.5mg/mL concentration) was applied to glow-discharged graphene-covered Quantifoil R1.2/1.3 copper/gold 300 mesh grids. The samples were preincubated on the grid for ~2 min within 100% humidified and 4˚C chilled Vitrobot Mark IV and plunged frozen in liquified ethane after blotting for 3-5s at blot force of -2. The grids were clipped and stored under a liquid nitrogen setting till screening and data collection.

***Single-particle cryo-EM data collection, processing, and analysis***

The grids were screened at ThermoFisher Scientific 200kV Arctica. The data collections were conducted at ThermoFisher Scientific 300kV Titan Krios equipped with Gatan K2 detector in EPU counting mode at a pixel size of 1.1 Å. More data collection details are listed in Table S1.

The movies for all datasets were processed in Cryosparc v3.3.1 (*5*). The workflow started with preprocessing with Patch Motion Correction and CTF estimation. The preprocessed micrographs were manually analyzed and selected for those with CTF estimated resolution less than 4 Å. A set of 2D references was generated using nsP1 map (EMDB ID: EMD-30796) (*1*) for the template picker for particle picking. The extracted particles were subjected to 2-3 rounds of 2D classifications to pick the best set of 2D classes estimated at 7 Å or less for *ab-initio* reconstructions first if necessary and followed by subsequent combinations of heterogenous refinements (2-3 classes) and 3D classifications (5-10 classes). The best 3D volume was selected for non-uniform refinement and uniform refinement at no symmetry (C1) applied with ctf and defocus optimizations and Ewald Sphere corrections (*5, 6*). These maps were refined to the range of 2.4-2.8 Å and reasonable Guinier plot Bfactor value for evaluations in both Cryopsarc and UCSF ChimeraX 1.3 (*7*). The PDB structures of nsP1, nsP2h, and nsP4 (7DOP, 6JIM, 7VB4, 7F0S) (*1-3*) were guided/fitted into the refined maps with models generated from DeepTracer (*8*) and AlphaFold2 (*9, 10*). The fitted structures were next exported to Coot (WinCoot 0.9.7 or Coot 0.9.5) (*11*) for chain morphing/refinement and manual building of protein chains and ligands. The built structures were refined in real-space refinement mode in Phenix (version 1.19.2) (*12*) to generate Table S1 to summarize all refinement statistics. The graphical illustrations and molecular analyses were prepared in ChimeraX (*13, 14*). All depictions of hydrogen bonds in less than 3 Å bond length were considered while the other molecular contacts were considered within ~4 Å distance. Multiple sequence alignment was conducted on MUSCLE server (*15*) and present as Weblogo (*16*) sequence conservation.

**Cellular cryo-ET sample preparation**

Human bone epithelial cell line, U2OS cells grown on fibronectin-coated gold 200 mesh R2/2 grids (Quantifoil) were infected with CHIKV-181 at an MOI of 50 for an incubation period of 8 hrs. The grids were then washed with PBS and a solution of 10 nm BSA gold tracer (Cat. #25486, EMS) was added directly before vitrification. Grids were blotted and plunged into liquid ethane using the LEICA EMGP plunge freezer device. Grids were stored under liquid nitrogen conditions until TEM data collection.

**Cryo-ET data collection and 3D reconstruction**

Grids of vitrified virus-infected cells were imaged on a Talos Arctica (ThermoFisher) operated at 200kV with a post-column energy filter (20eV) and K2 Summit detector with a calibrated pixel size of 3.54Å. Single-axis, bi-directional tilt series were collected using SerialEM software with low-dose settings and defocus range of -3 to -5.5 µm. The total average dose at the specimen was $90e^{-}/A^{2}$, distributed over 51 tilt images, covering an angular range of -50° to +50° with an angular increment of 2°. Motion between frames of each tilt image in the tilt series was corrected using patch-tracking in MotionCor2 software (*17*). Tilt images were compiled, automatically aligned and reconstructed into 3D tomograms using EMAN2 software (*18, 19*). In total, 146 tomograms were judged as sufficient quality for further subvolume analysis.

**Subvolume averaging and analysis**

To generate an initial model of the RC at the base of RNA spherules, 50 high SNR particles in different orientations were picked from tomograms at x4 binning. These RC particles were input to the EMAN2 initial model generation program, performed in two steps. First, particles were iteratively aligned with C1 symmetry for three iterations and aligned to the symmetry axis. Then an additional five iterations were performed, applying C12 symmetry to produce an initial model with low-resolution features of the spherule base, plasma membrane, and RC. This initial model was filtered to 40Å resolution and used for subsequent 3D subtomogram refinement of the full dataset. Subtomogram averaging was then performed with 724 RC particles at x1 binning (3.54Å/pix) from 49 tomograms while applying C12 symmetry with a loose mask around RC protein density to exclude the surrounding membrane from alignment. After subtomogram refinement and additional sub-tilt refinement of particle translation and rotation produced a final RC map at 7.3Å resolution (Fig. 3D, Fig. S6). To determine a C1 reconstruction of the RC, the 12 possible asymmetric orientations based on the previous refinement with C12 symmetry were iteratively searched using the *breaksym* option in EMAN2 subtomogram refinement. After 10 iterations this resulted in a converged map with asymmetric density features in the central region of the RC shown in Fig. 3E.

In a similar way to RCs, 100 high SNR particles appearing as small rings on the inner leaflet of the plasma membrane were manually picked from tomograms and subjected to EMAN2 initial model generation at x4 binning. After three iterations of C1 alignment, with the result aligned to the symmetry axis, and five additional iterations applying C12 symmetry, a low-resolution map displayed a ring-like structure docked to a membrane. This map was again blurred to 40Å resolution and used as the input for five iterations of subtomogram refinement of 5704 ring particles at x1 binning (3.54 Å/pix) with C12 symmetry applied. A soft mask was applied to focus alignment of particles on the protein ring density rather than the membrane. Subtomogram refinement and sub-tilt refinement of translation and rotation, again with C12 symmetry applied, produced a final map at 9.9Å resolution (Fig. 4C, Fig. S6). After 10 iterations of symmetry-breaking refinement, a C1 map of the ring revealed asymmetric density within the central pore, shown in Fig. 4D.

Refined ring particle orientations determined by subtomogram averaging were mapped to the originating tomograms using EMAN2 tool e2spt_*mapptclstotomo.py*. Visualization and model docking was performed using UCSF ChimeraX and its built-in fit-in-map tool (*13, 14*).

**Figure S1 Cryo-EM analysis of nsP1+2+4 RC core.** (A) cryo-EM workflow in Cryosparc at C1 refinement symmetry. (B) FSC resolutions of the map: masked versus unmasked. (C) map resolution overview and the overall geometrical position at the spherule. Scale bar (blue-red: 2.5-4.5 Å) (D) Viewing the direction distribution plot of the images to show the precision at the map posterior view.

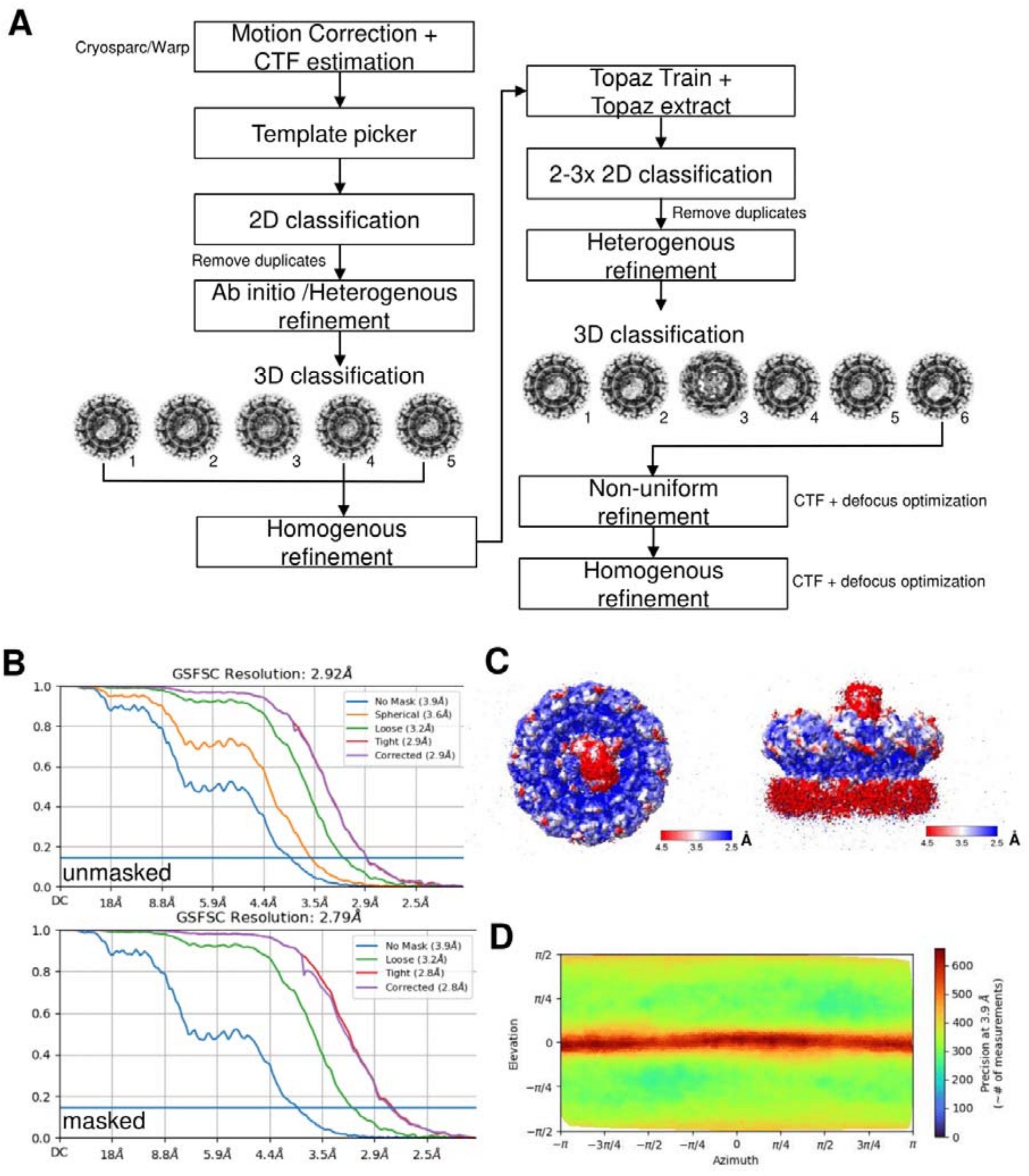

**Figure S2. Map density fitting quality at the interaction interfaces of the RC core and nsP4 sites.** The density map fitting (black mesh) on focused regions of (A) nsP2-4 interface, (B) nsP2h-RNA interface, (C) nsP1-4 interface at nsP1 chain L site, and (D) GDD in the nsP4 palm active site (also known as motif C) and its first tyrosine residue on nsP4 N-terminus tip. All parts are colored according to main figures and supplementary figures.

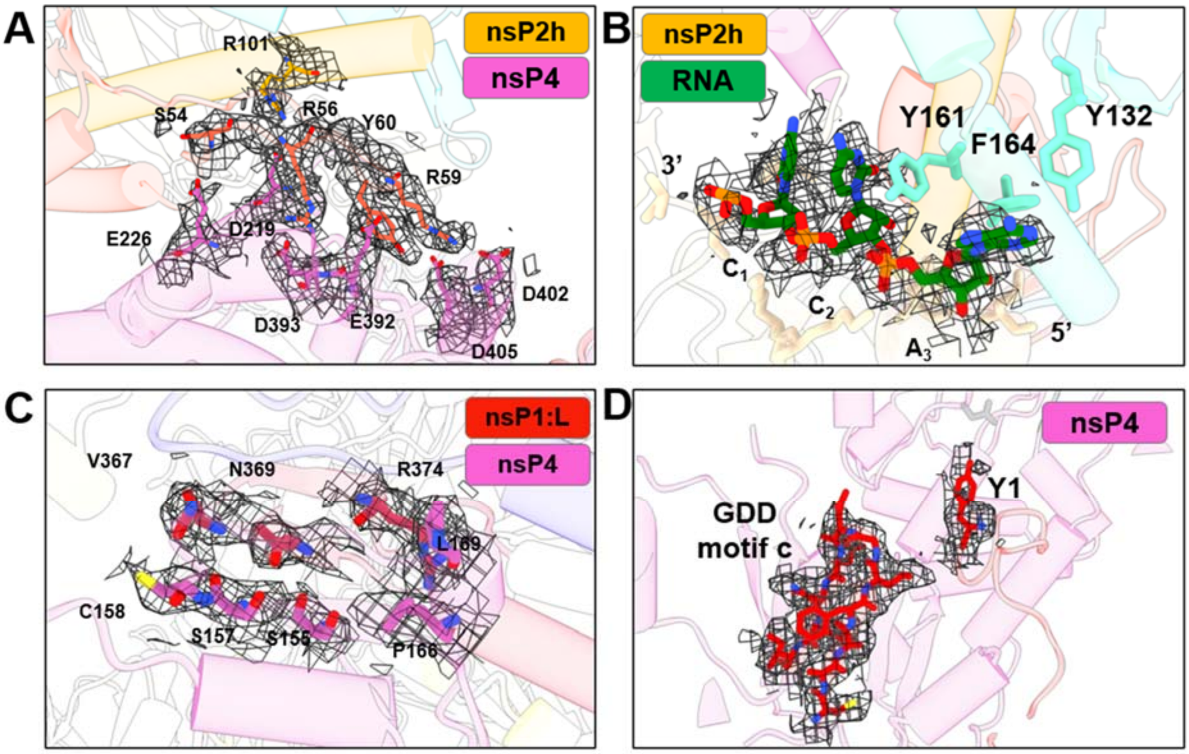

**Figure S3. The structural interfaces in the RC and their sequence conservation.** (A) The interaction of nsPs (nsP1: A-L; nsP4: X; nsP2-RNA: Y-Z) are presented here with their chain name in the form of network linking nodes and individual interface area. (B) The subdomain layouts of nsP2, nsP4 and nsP1 show their interacting interfaces via connections by black line. (C-F) The Weblogo (*20*) style sequence conservations of the interacting residues are compared through multiple-sequence-alignment (MSA) among the alphaviruses on their nsPs. The interface residues of nsP2h-nsP4 are shown in boxes for nsP2h part in (C) while for nsP4 in (D). The interface residues which are highlighted here in boxes are for nsP4 in (E) and for nsP1 in (F). The sequences are colored based on charges (positive blue; negative red). [Alphaviruses were used in multiple-sequence alignment (MSA): chikungunya virus (CHIKV; NC_004162), o'nyong nyong virus (ONNV; AF079456.1), Semliki forest virus (SFV; NC_003215.1), Sindbis virus (SINV; NC_001547.1), Ross River virus (RRV; GQ433354.1), Mayaro virus (MAYV; NC_003417.1), Barmah forest virus (BFV; MN689034.1), Venezuelan equine encephalitis virus (VEEV; L01442.2); Eastern equine encephalitis virus (EEEV; EF151502.1), Western equine encephalitis virus (WEEV; MN477208.1); Eilat virus (EILV; NC_018615.1), and Getah virus (GETV; NC_006558.1).] interaction type: hydrophilic (star), hydrophobic (circle) work in progress to add in hydrophobic int.

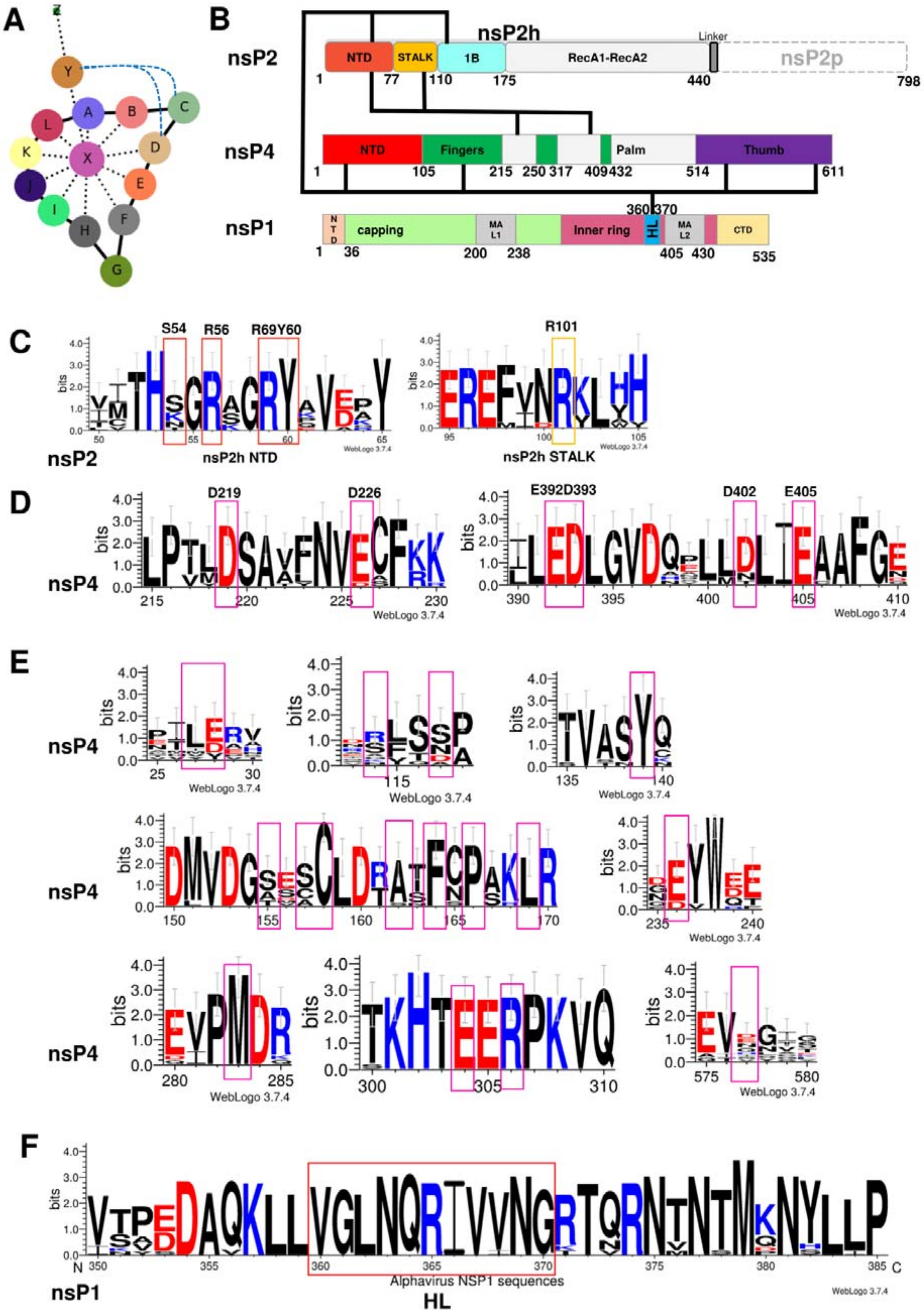

A
B
nsP2
nsP2h
NTD
STALK
1B
RecA1-RecA2
Linker
nsP2p
nsP4
Fingers
Palm
Thumb
nsP1
capping
MA L1
Inner ring
HL
MA L2
CTD
C
S54
R56
R69Y60
R101
nsP2h NTD
nsP2h STALK
D
D219
D226
E392D393
D402
E405
E
F
Alphavirus NSP1 sequences

1

**Figure S4. Comparison between cryoEM structure nsP4 from ONNV and crystal structure nsP4 from SINV and representative viral RdRp from EV71.** (A) the cryoEM structure of nsP4 from ONNV was colored according to the subdomain (NTD: red; Index finger: green; Middle, ring, and pinky fingers: orange; Thumb: purple). (B) the crystal structures of the elongation complex from *Picornaviridae*: enterovirus-71 (EV71) RdRp with its RNA substrate (PDB 7KWQ) (RNA colored blue). (C-D) The crystal structrures of alphavirus nsP4 homologues: RRV and SINV. (E) The crystal structure of another homolog from *Flaviviridae*: classical swine fever virus RdRp (CSFV; PDB 5Y6R). ONNV has r.m.s. deviation (RMSD) of 1.0-1.2 Å (194 Cα aligned in aa215-575) at RdRp core domain when superimposed to nsP4 from RRV and SINV. The rearrangement of the index finger (black dotted arrow) is shown in (D) to form the folding in (A). (A-B) When compared to EV71 RdRp (PDB 6KWQ), it has 1.3 Å (57 Cα aligned) at the RdRp core domain. The RMSD between ONNV nsP4 and CSFV RdRp is 1.26 Å.

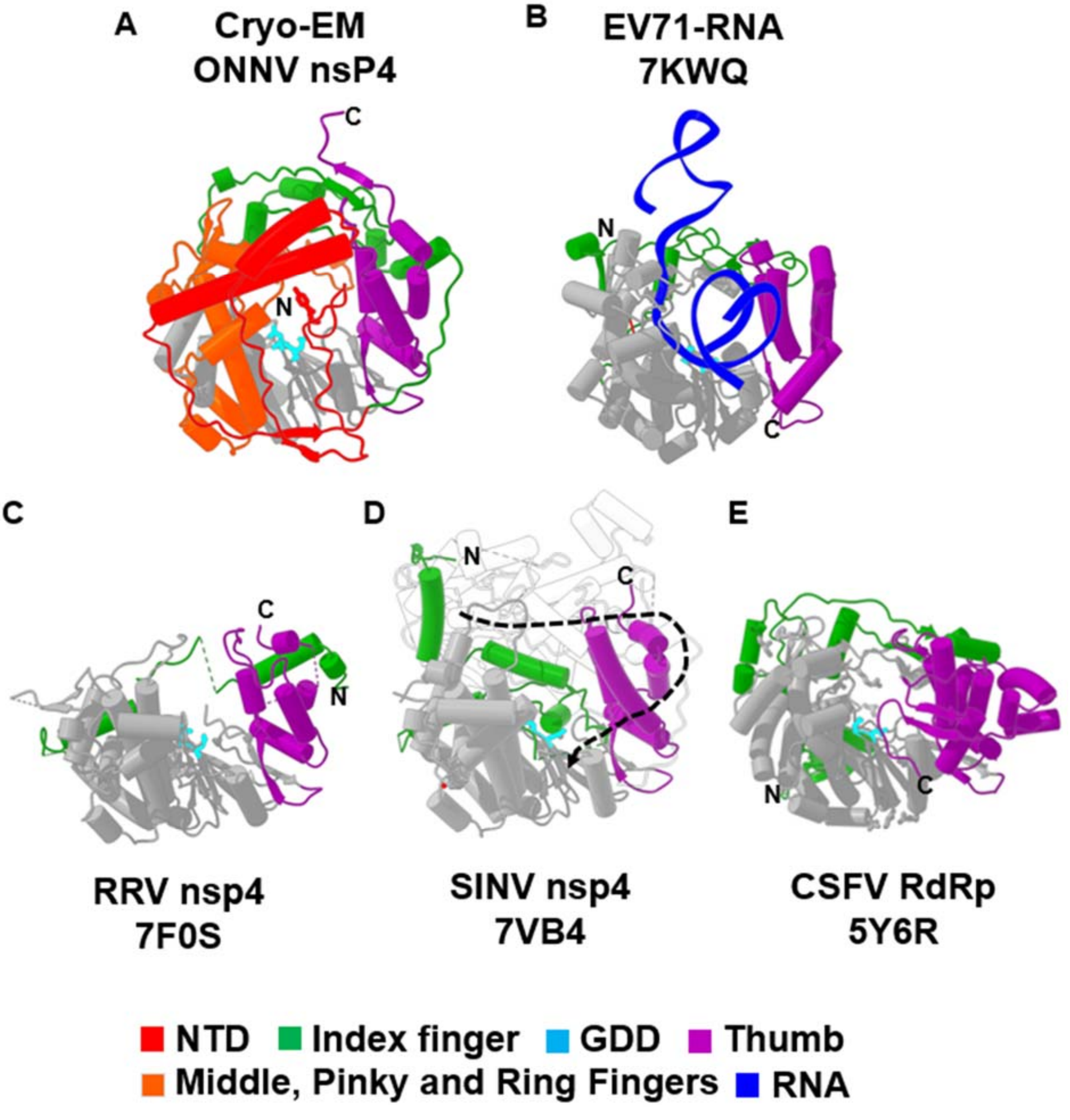

**Figure S5. Subtomogram average map analysis of RC and non-replicative nsp complex.** (A) RC density map shown as transparent surface with nsp1+4 atomic model (rainbow) rigid fit into density. Single nsp1 subunit is computationally extracted (dark grey) and backbone of nsp1 subunit model is displayed (rainbow). (B) FSC curves of the masked and unmasked maps. (C) RC map colored by local resolution. (D) Non-replicative nsp ring density map displayed as transparent surface with nsp1+4 atomic model rigid fit into density, with single nsp1 subunit displayed. (E) Gold-standard FSC curve of masked and unmasked map with (F) density map colored by local resolution.

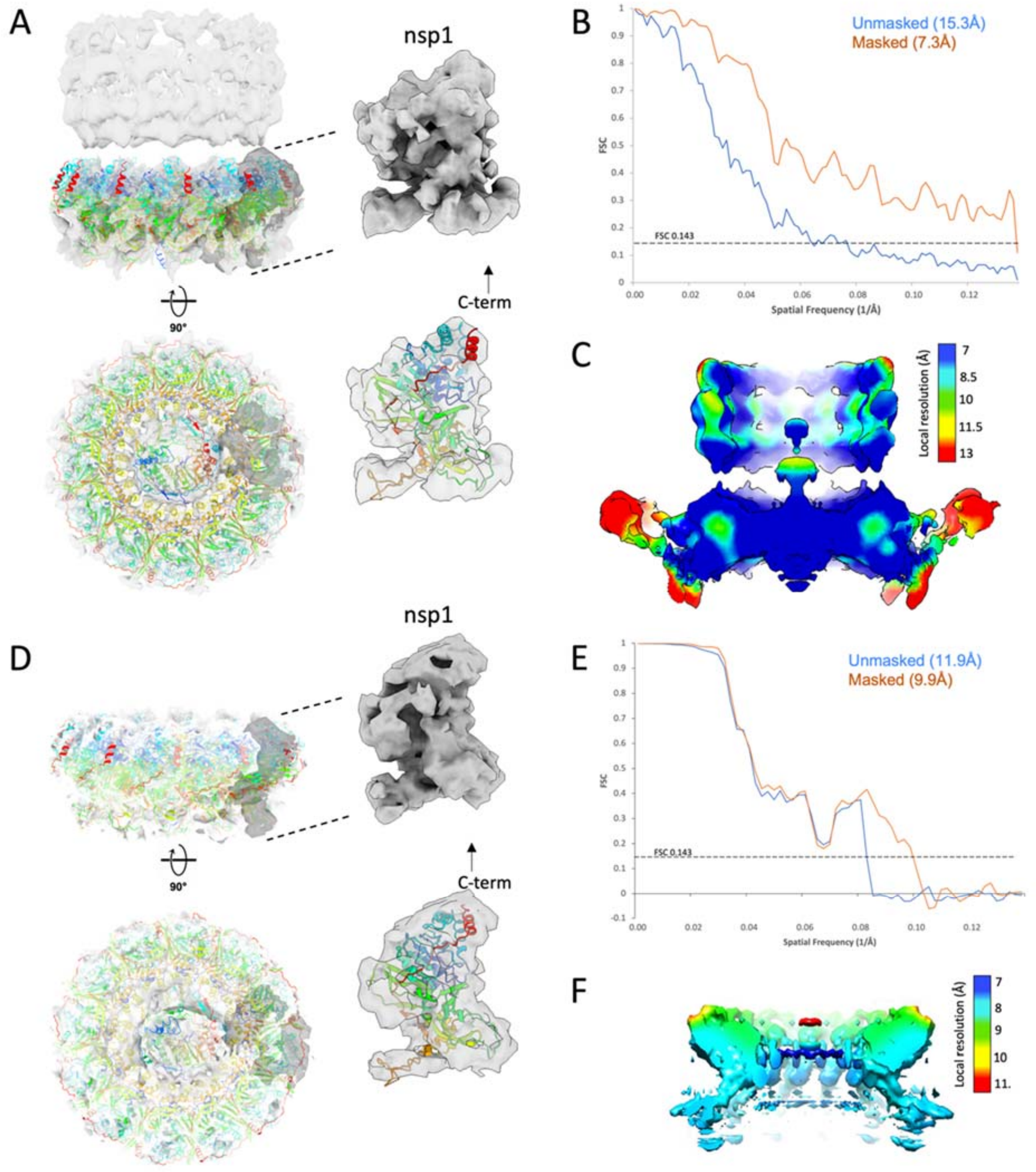

**Figure S6. Non-replicative nsp complexes on the plasma membrane.** (A) Low-mag montage of the cell periphery reveals numerous cell extensions (red arrows) emanating outward from the cell body. Scale bar 2 micron. (B) Slice image of a tomogram reconstruction displaying a representative thin cell extension. Zoom-in slice views of different membrane surfaces containing nsp complexes (white arrows). (C) Asymmetric 3D class averages of nsp complexes reveals presence of nsP2+4 in central pore of nsP1 ring in each class.

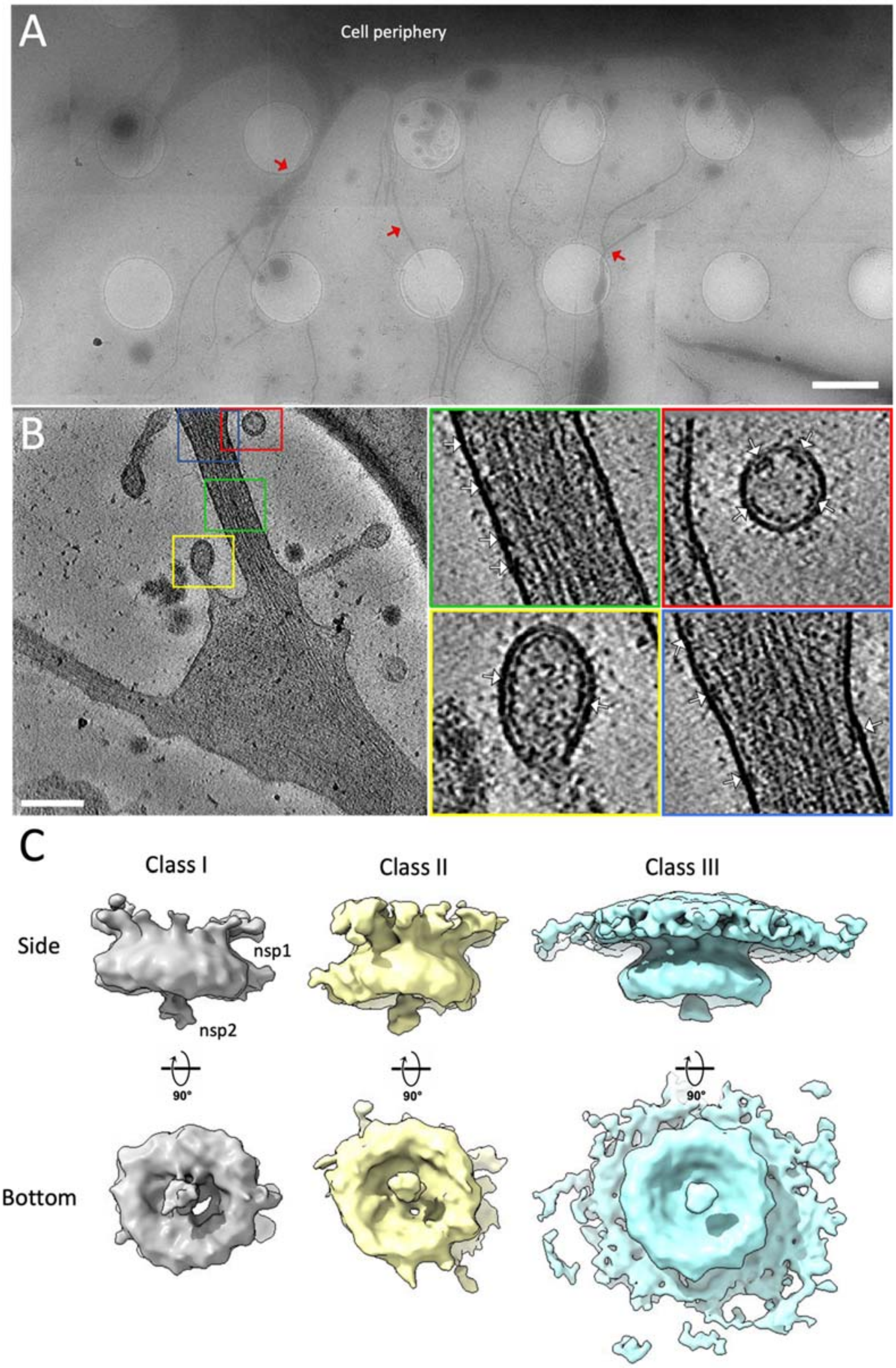

**Figure S7. Membrane association of the active RC via the outer surface of the nsP1 ring.** MA Loops 1 and 2 have been reported previously (*1, 21*). The MA patches include mainly hydrophobic and positively charged residues 125-126, 269-270, 451-455, and 457-470 which collectively form a surface belt on the nsP1 upper ring.

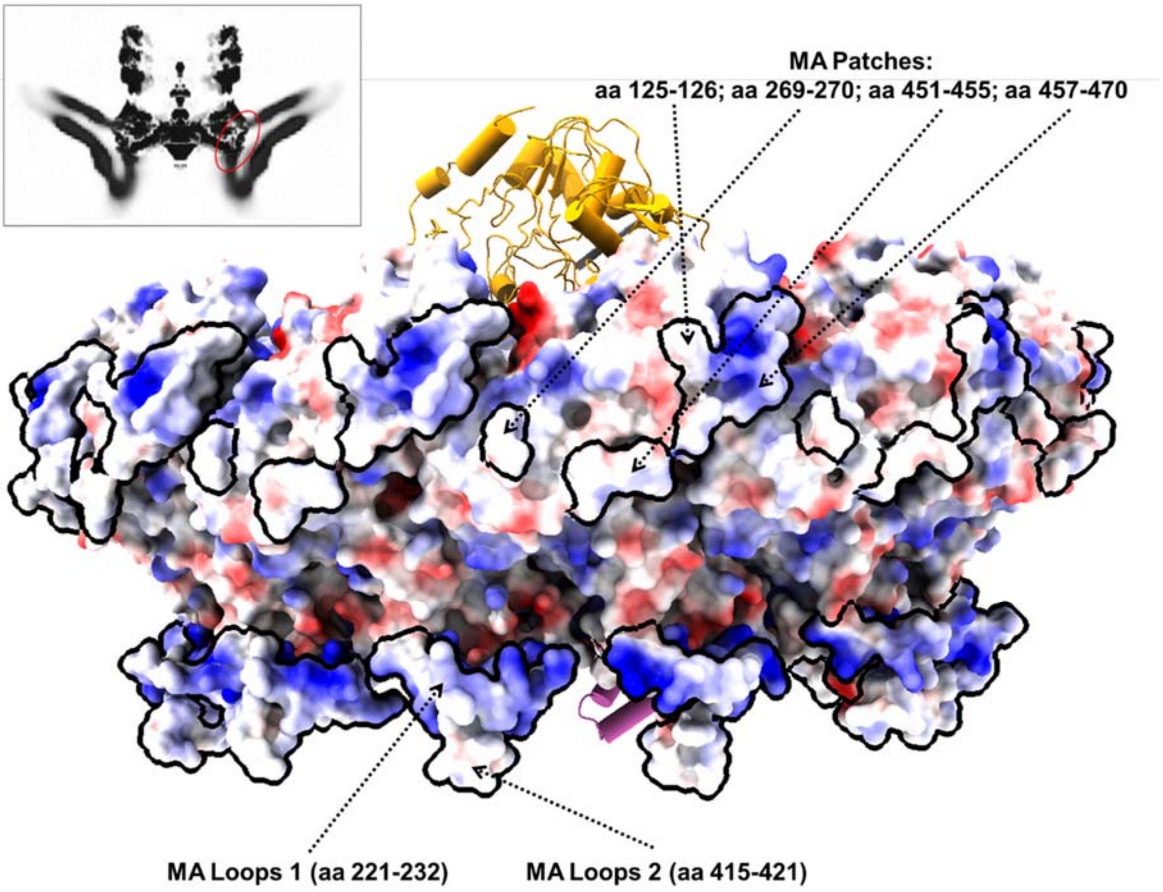

**Figure S8. Topology of the (+) RNA virus replication complexes.** Left: MHV nsp3 pore at the neck of the double-membrane vesicles (DMVs) derived from ER membrane (*22*). Middle: CHIKV replication complex at the neck of the spherule derived from the plasma membrane. Right: FHV protein A crown at the neck of the viral spherue derived from the mitochondrial outer membrane (*23*).

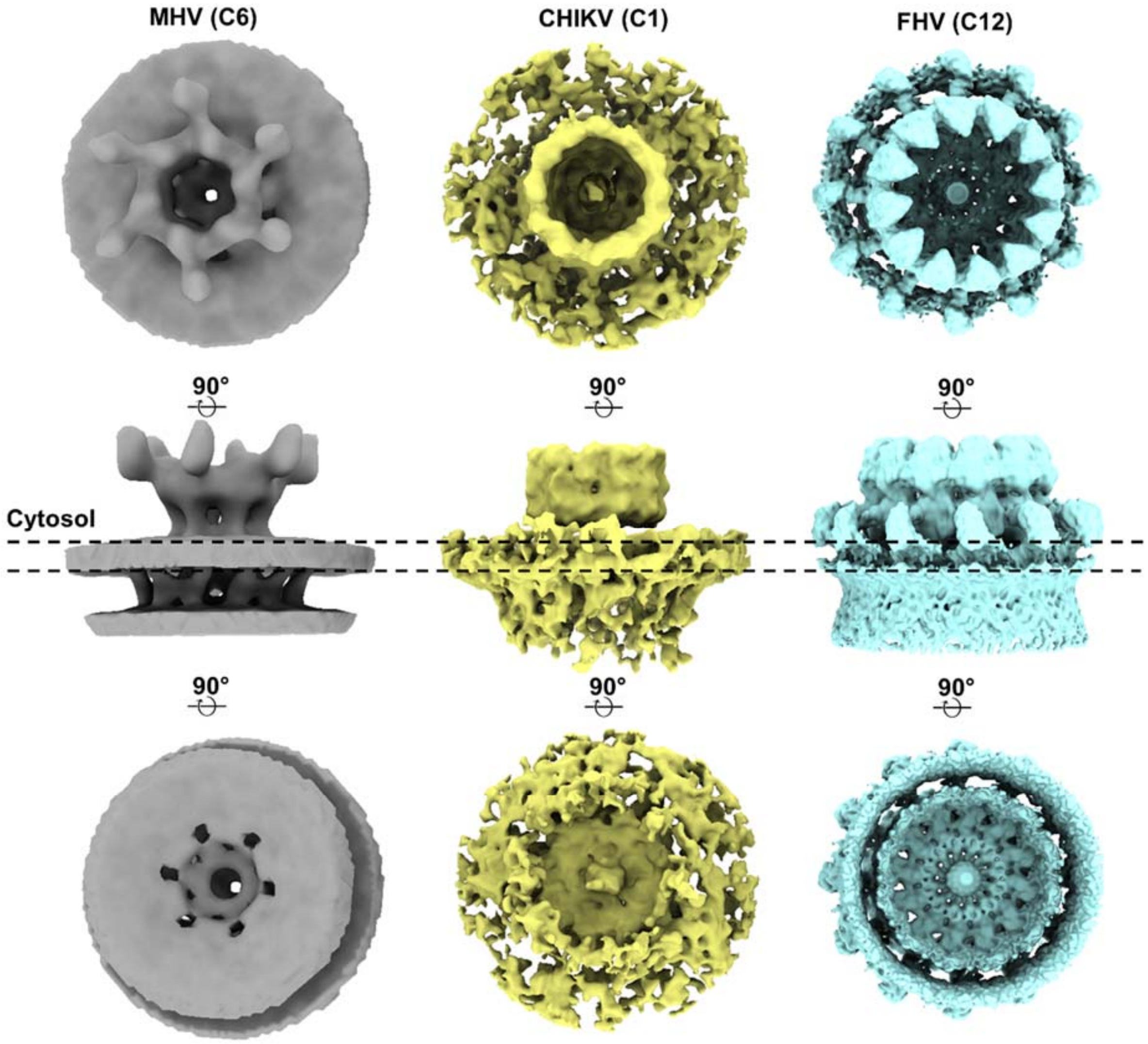

**Table S1**

| | P124 |
|---|---|
| | EMDB: |
| | PDB: |
| **Data collection and processing** | |
| Detector | Gatan K2 |
| Magnification | 130,000 × |
| Voltage (kV) | 300 |
| Electron exposure (e–/Å$^2$) | 34 |
| Defocus range (μm) | -0.5 ~-3.2 |
| Pixel size (Å) | 1.10 |
| Symmetry imposed | C1 |
| Initial particle images (no.) | 898,911 |
| Final particle images (no.) | 132,510 |
| Map resolution (Å) | 2.8 |
| FSC threshold micrographs | 0.143 |
| Map resolution range (Å) | 2.4-6.2 |
| **Refinement** | |
| Initial model used (PDB code) | 7DOP |
| Model resolution (Å) | 2.8 |
| FSC threshold | 0.143-0.5 |
| Model resolution range (Å) | 2.8~3.3 |
| Map sharpening B-factor (Å$^2$) | ~63 |
| **Model composition** | |
| Non-hydrogen atoms | 53,139 |
| Protein residues | Protein: 6704; |
| Nucleotide | 3 |
| Water | 0 |
| Ligands | ZN: 12 |
| | GTP: 12 |
| | ATP: 12 |
| B factors (Å$^2$) | |
| Protein | 24.37/215.98/81.44 |
| Nucleotide | 215.58/231.42/224.21 |
| Water | --- |
| Ligand | 56.25/126.14/65.90 |
| **R.m.s. deviations** | |
| Bonds length (Å) | 0.004 |
| Bonds Angle (˚) | 0.66 |
| **Validation** | |
| MolProbity score | 1.59 |
| Clashscore | 3.83 |
| Q-score | |
| Rotamer outliers (%) | 0.05 |
| **Ramachandran plot** | |
| Favored (%) | 93.72 |
| Allowed (%) | 6.15 |
| Disallowed (%) | 0.14 |
